# Refined Myocardium Segmentation from CT Using a Hybrid-Fusion Transformer

**DOI:** 10.1101/2024.09.27.615510

**Authors:** Shihua Qin, Fangxu Xing, Jihoon Cho, Jinah Park, Xiaofeng Liu, Amir Rouhollahi, Elias J. Bou Farhat, Hoda Javadikasgari, Ashraf Sabe, Farhad R. Nezami, Jonghye Woo, Iman Aganj

## Abstract

Accurate segmentation of the left ventricle (LV) in cardiac CT images is crucial for assessing ventricular function and diagnosing cardiovascular diseases. Creating a sufficiently large training set with accurate manual labels of LV can be cumbersome. More efficient semi-automatic segmentation, however, often includes unwanted structures, such as papillary muscles, due to low contrast between the LV wall and surrounding tissues. This study introduces a two-input-channel method within a Hybrid-Fusion Transformer deep-learning framework to produce refined LV labels from a combination of CT images and semi-automatic rough labels, effectively removing papillary muscles. By leveraging the efficiency of semi-automatic LV segmentation, we train an automatic refined segmentation model on a small set of images with both refined manual and rough semi-automatic labels. Evaluated through quantitative cross-validation, our method outperformed models that used only either CT images or rough masks as input.

## 2. Introduction

Early detection of morphological and functional abnormalities of the myocardium is essential for preventing cardiovascular diseases [1], which remain a leading cause of mortality worldwide [2]. In cardiology, ventricular performance is typically assessed through measurements such as the wall thickness, ventricular volume, and ejection fraction [3]. These critical indices are often obtained through the segmentation of cardiac images produced by medical imaging modalities such as magnetic resonance imaging and computed tomography (CT) [4].

Recent advancements in deep neural networks have greatly improved medical image segmentation [5]. Convolutional neural network (CNN)-based learning models, such as the U-Net [6], are commonly adopted in segmentation [7], [8], [9]. These models, however, typically require large manually labeled datasets, which are time-consuming and costly to produce, especially for 3D volumes. Consequently, cardiologists often use semi-automatic labeling to prepare training data [10], [11]. Unfortunately, such intensity-based methods struggle with fine details, such as separating the left ventricle (LV) wall from adjacent papillary muscles in CT images [12]. Several previous studies have shown that the exclusion of papillary muscles yields the most reproducible endocardial segmentation [13], [14]. Additionally, a smooth endocardium boundary can ensure more accurate measurements of the ejection fraction and the volume [15]. Thus, proper refinement of the semi-automatic coarse masks – which cannot be practically achieved by resource-demanding manual delineations – is essential for creating papillary-muscle-free input labels for deep-network training.

In this work, we extend the Hybrid-Fusion Transformer (HFTrans) segmentation model [16] to accommodate two input channels for refining the segmentation and removing the papillary muscles from CT images of the myocardium. Trained on a small dataset (representing the scarce manually labeled set), the model takes both CT images and semi-automatically created coarse masks as two-channel input, learning to produce refined masks. The CNN performs coarse segmentation, while the transformer refines the masks to exclude papillary muscles [17]. By leveraging minimal fine labels, our model efficiently enhances the practicality of semi-automatic labeling, reducing the need for extensive manual refinement. Evaluation shows improved segmentation accuracy of the two-channel (over single-channel) input.

## 3. Materials and Methods

### 3.1 Data Description

We retrospectively identified a dataset of 35 adult patients with severe aortic stenosis who underwent transcatheter aortic valve replacement (TAVR) from 2018 to 2020. These patients were selected from a pool of 272 subjects that we have previously reported [18] and used to develop a pipeline for digital twin reconstruction of the aorta, aortic root, and aortic valve, allowing for a more comprehensive analysis of valve calcification burden and distribution [7]. (In the current study, however, we apply an HFTrans-based framework to refine LV segmentation in cardiac CT images by excluding papillary muscles.) We excluded patients under 18 years of age, as well as those with mixed aortic regurgitation, more than mild concomitant valvular disease, a history of ischemic heart disease, prior valve or coronary artery bypass graft surgery, or poor-quality pre-TAVR gated cardiac CT images. Cardiac CT images were retrospectively collected in the DICOM format with approval from the local institutional review board [19]. The raw DICOM data were initially de-identified by an independent party, who converted the images into the nearly raw raster data (NRRD) format. Images were downsampled to the size 160×160×160, resulting in an in-plane pixel size varying from 0.89 to 1.44 mm, and a slice thickness varying from 0.69 to 1.15 mm.

A trained expert utilized 3D Slicer [20] to segment the LV semi-automatically with the “grow from seed” option, a region-growing method that classifies myocardial regions using seeds for foreground (LV) and background. Misclassified regions were iteratively corrected by adding seeds until the desired segmentation was achieved, then converted into a binary label file. To exclude the papillary muscles, five cases containing papillary muscles were selected. A clinician manually edited each slice to remove the papillary muscles using the erasing tool in 3D Slicer.

### 3.2 Methods

For each training CT image, *I*, we denote its semi-automatic *coarse* LV mask containing papillary muscles as *M*^*c*^ and its manually *refined* mask without the papillary muscles as *M*^*r*^. The proposed network focuses on leveraging information from both *I* and *M*^*c*^ to predict a refined, smooth LV region, given a limited availability of cases with *M*^*r*^.

The deep-learning model we propose to use is based on the HFTrans architecture [16]. HFTrans combines the strengths of both CNNs and transformers, where the CNN encoder is used to extract contextual features from each input image-label pair, and the transformer layers exert self-attention on these extracted features to enhance representation and connect information across channels. Based on this framework, we propose to expand the input image to two channels defined in ℝ^2×*W* ×*H* ×*D*^ (channel, width, height, depth). For the *i*^*th*^ training image, Channel 1 is the original cardiac CT volume *I*_*i*_ ∈ ℝ^1× *W* × *H* × *D*^ and Channel 2 is its corresponding coarse mask 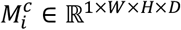. The stacked new input 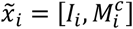 provides a richer context, where the coarse mask captures the overall structure of the LV (where the refined mask should be confined to) and its surrounding anatomy, and the CT image provides voxel-level details. Correspondingly, the manually refined mask 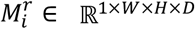 serves as the training label for the *i*^th^ image. This augmented input enables the transformer part of the network to focus on correcting regional details and producing a refined mask.

For each input pair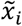, the features *f*_*j*_ (where *j* = 1, 2, 3) are extracted from three CNN encoders. The first encoder performs early fusion by processing both the CT image and rough mask together, while the other two encoders extract features individually from each sequence (*I*_*i*_ *or* 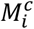). These features are then linearly projected, and a positional embedding is added to retain spatial information, as shown in Figure 1. The initial embedding *z*_*o*_ is calculated as:

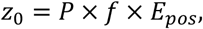

where *P* is the projection matrix and *E*_*pos*_ is the positional embedding. The transformer then applies a multi-head self-attention (MSA) and a multilayer perceptron (MLP), with layer normalization (LN), to refine the features across multiple layers, with each Transformer layer defined as:

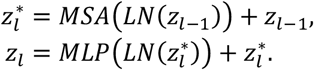

**Figure 1.**
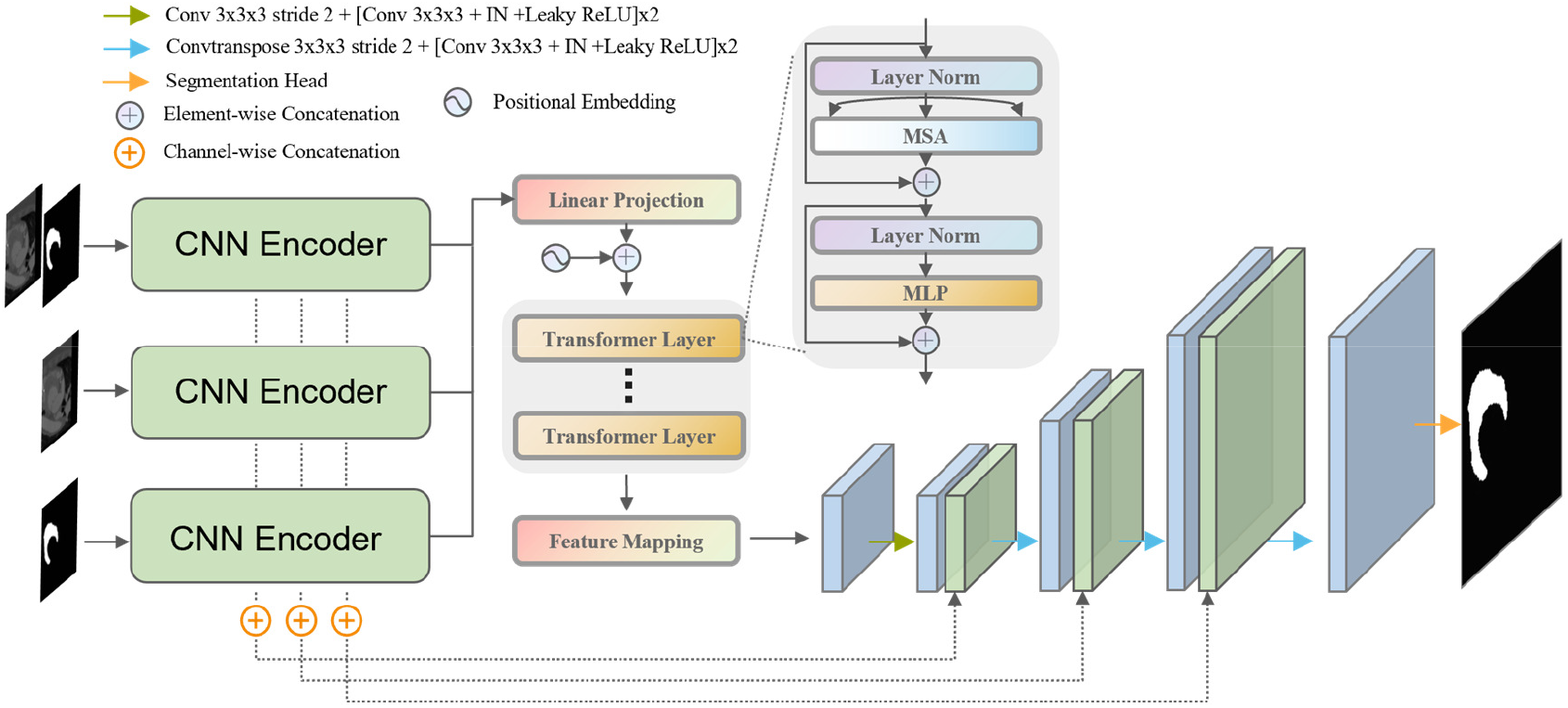
Overview of our neural network structure. HFTrans consists of three CNN encoders. The first encoder performs early fusion of the CT images and the rough mask, while the other two encoders are used to extract contextual feature representations from the CT image and rough mask separately. A Transformer encoder is then used to integrate the features extracted by these encoders. CNN=convolutional neural network, Conv=convolutional layer, IN=Instance Normalization, Layer Norm=layer normalization, MSA=multi-head self-attention, MLP=multilayer perceptron.

To deal with limited input data with refined labels, we employed several data augmentation techniques, including linear transformations such as random rotations (−30° to 30°) and isotropic scaling (0.7–1.4), on both the CT images and their corresponding masks, while explicitly excluding shear. In addition, we applied intensity augmentations, including Gaussian noise, Gaussian blur, random brightness, contrast, and gamma adjustments, as well as random flipping along the three spatial dimensions. Due to GPU memory constraints and for further augmentation, the original input size of 160×160×160 was cropped to 128×128×128 at a random location. During the pre-processing stage, we applied contrast adjustments, added Gaussian noise, and performed histogram equalization on the CT images. For post-processing, we enhanced model stability and performance by flipping the input patch along all three dimensions, generating 8 different spatial perspectives of the same object. We then predicted 8 probability maps, and after flipping them back to their original orientations, we averaged them to create the final probability map, improving performance and stability.

### 3.3. Validation

We employed a leave-one-out cross-validation strategy to maximize the utility of our dataset, where four cases were used for training and the remaining case was reserved for validation. Augmentation and cropping strategies were explicitly constrained to avoid overlap between training and validation sets. Each model was executed using PyTorch (https://pytorch.org/version1.13.1) with parameters initialized randomly. This approach ensures that every data point is used both for training and validation in different experiments, providing a robust estimate of the model’s performance.

We adopted two key metrics: the Dice similarity coefficient (DSC) and the 95% Hausdorff Distance (HD95) [21]. The DSC evaluates the overlap between the predicted and ground-truth masks, providing a measure of segmentation accuracy. The HD95, on the other hand, assesses the boundary alignment, ensuring that the refined segmentation is not only accurate but also spatially consistent with the ground truth.

To determine the contribution of each channel of 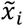 on the performance of the model, we designed and tested three variations: using the original CT image 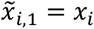, using the rough mask alone 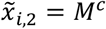, and using the combined stack of the CT image and rough mask 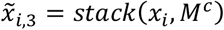. Cross-validation was employed for each type of input data. The differences in the DSC and HD95 were evaluated using paired *t*-tests to determine the mean differences. These statistical analyses were performed utilizing the R stats package, version 4.42 (https://cran.rstudio.com/bin/windows/base/NEWS.R-4.4.2.html). A *p*-value of less than 0.05 was considered indicative of statistical significance.

## 4. Results

The training took an average of three hours for 5000 epochs using an NVIDIA RTX A6000 GPU with 48 GB of memory, on an Intel Xeon Gold 6342 2.8GHz 24 Core CPU with 100 GB of RAM.

### 4.1. Quantitative results

The results are presented in Tables 1 and 2, as well as Figure 2. When using only CT images as input, the model struggled to achieve satisfactory performance, likely due to limited training data. The average DSC was 87.2%, with a high standard deviation for both DSC (6.7) and HD95 (12.20). In contrast, the two-channel input approach ([*CT,M*^*c*^]) achieved the improved mean DSC of 95.2% and an average HD95 of 2.12, with considerably lower variability, as reflected by the standard deviations of DSC (1.6) and HD95 (0.71).

**Table 1:** Comparison of three different input strategies.

| | $CT$ | | $M^c$ | | $[CT, M^c]$ | |
| --- | --- | --- | --- | --- | --- | --- |
|  | DSC↑<br>(%) | HD95↓<br>(voxels) | DSC↑<br>(%) | HD95↓<br>(voxels) | DSC↑<br>(%) | HD95↓<br>(voxels) |
| Case 1 | 90.4 | 2.78 | 94.7 | 2.58 | **96.8** | **1.41** |
| Case 5 | 87.4 | 31.76 | 89.9 | 5.67 | **96.2** | **1.72** |
| Case 16 | 93.3 | 5.76 | **95.9** | 2.18 | 95.5 | **2.00** |
| Case 19 | 75.8 | 8.68 | **97.0** | **1.31** | 92.7 | 3.27 |
| Case 34 | 89.1 | 2.91 | 92.3 | 3.42 | **94.6** | **2.21** |
| Mean | 87.2 | 10.38 | 94.0 | 3.03 | **95.2** | **2.12** |
| StD | 6.7 | 12.20 | 2.9 | 1.66 | **1.6** | **0.71** |

Note. - $CT$ =CT image only input, $M^c$ =coarse mask only input, $[CT, M^c]$ =combined stack of the CT image and rough mask input, DSC=Dice similarity coefficient, HD95=95% Hausdorff Distance, Mean=mean value of all cases, StD=standard deviation.

**Table 2:** Results of two-sided paired *t*-tests for mean differences in DSC and HD95 between three distinct input strategies.

|  | DSC (%) |  |  |  | HD95 (voxels) |  |  |  |
| --- | --- | --- | --- | --- | --- | --- | --- | --- |
|  | Mean<br>Difference | t | p | CI95 | Mean<br>Difference | t | p | CI95 |
| $[CT, M^c] - CT$ | 8.0* | 3.2 | 0.03 | [1.13, 14.80] | -8.3 | -1.50 | 0.21 | [-23.56, 7.05] |
| $M^c - CT$ | 6.8 | 1.9 | 0.14 | [-2.83, 16.46] | -7.4 | -1.50 | 0.21 | [-23.90, 8.17] |
| $[CT, M^c] - M^c$ | 1.2 | 0.7 | 0.53 | [-2.21, 4.50] | -0.9 | -0.95 | 0.39 | [-2.02, 1.24] |

Note. - $CT$ =CT image only input, $M^c$ =coarse mask only input, $[CT, M^c]$ =combined stack of the CT image and rough mask input, DSC=Dice similarity coefficient, HD95=95% Hausdorff Distance, $t$ = $t$ -statistic, $p$ = $p$ -value, CI95=95% confidence interval.
\* Statistically significant at $p < 0.05$

**Figure 2.**
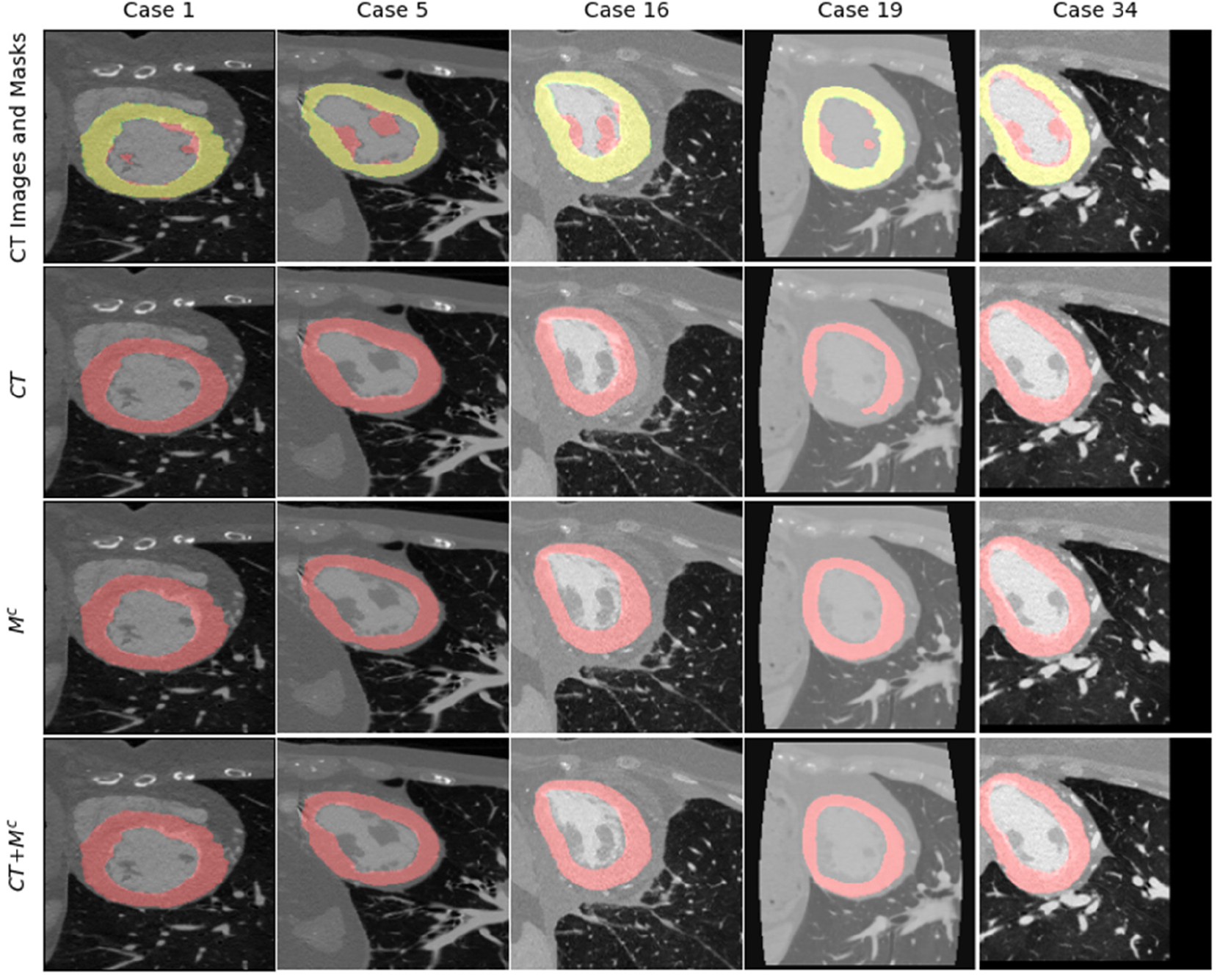
Qualitative comparison of different input types across the 5 cases. In the first row, the manual green mask represents the LV wall excluding the papillary muscle, while the semi-automatic red mask indicates the unwanted papillary muscle (yellow is the overlap of the two; green is hardly seen as the refined mask is a subset of the coarse mask). The remaining three rows show the model-predicted LV segmentation using different input types: CT image only (*CT*), coarse mask only (*M* ^*c*^), and the combined stack of the CT image and rough mask (*CT* + *M*^*c*^).

Table 2 presents the results of the paired *t*-tests examining the differences in DSC and HD95 across three distinct input strategies. The analysis indicates that the two-channel input method demonstrated a statistically significant (p = 0.03) enhancement in DSC, alongside a (non-significant) reduction of 8.3 in HD95 when juxtaposed with the CT-only input. In contrast, the disparity between the rough mask-only input strategy and the CT images-only input was less pronounced, yielding non-significant results for both DSC and HD95. Furthermore, the comparison between the combined approach and the rough mask-only method indicated a gain of 1.2% in DSC and a decrease of 0.9 voxels in HD95 (both non-significant).

### 4.2. Applicability

To qualitatively evaluate our method’s effectiveness in removing papillary muscles on more images, Figure 3 shows refined LV wall predictions for the remaining 30 subjects, using a two-channel model trained on the five manually refined cases. The model successfully excludes papillary muscles, demonstrating its practical value in scenarios where only a small refined training set is available.

**Figure 3.**
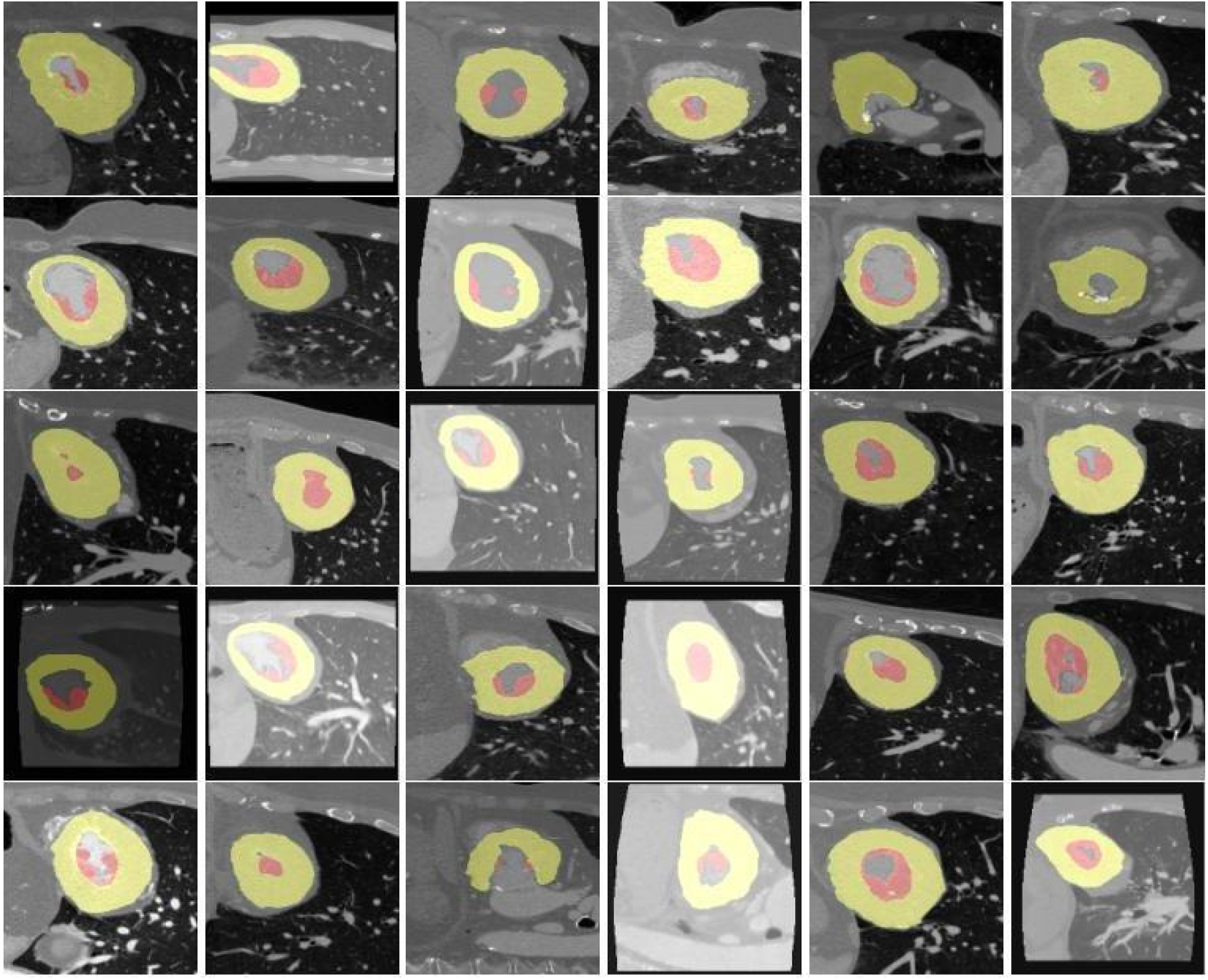
Refined segmentation predictions for LV walls of the remaining 30 adult patients. Each patient’s dataset includes a semi-automatically generated mask (including the papillary muscles) highlighted in red. Automatic segmentation is in green, with the intersection of the two masks in yellow. (Green is rarely seen, as the refined mask is expected to be included in the coarse mask.)

## 5. Discussion

In this paper, we presented a two-channel HFTrans-based deep-learning method for refining semi-automatic LV segmentation in cardiac CT images, with a focus on excluding the papillary muscles. By utilizing both the CT images and rough masks as input, the method integrates complementary information to enhance segmentation accuracy and stability, which is particularly useful in clinical scenarios when refined manual labels for training are scarce. Although the two-channel method did not outperform the single-channel rough-mask input in all cases, the inclusion of the CT image remains valuable. Indeed, it provides essential anatomical context around the LV, helping the model to refine segmentation where the rough mask alone may miss critical details. By combining the structural information from rough masks with the contextual data from CT images, the proposed method generally yields more reliable and consistent segmentation results in most cases.

We trained our model on a very small set of 5 subjects with refined labels. Our results, showing improvement with a two-channel input, serve as proof of principle that rough semi-automatic labels, when fed to a model trained on a small set of manually and laboriously created refined labels, can be efficiently utilized in automatic refined segmentation. It is noteworthy that, in segmentation model training, every voxel in every subject’s 3D image counts as a data point with a unique ground-truth label value. This creates ample training data, even if only a handful of subjects are included.

The non-significant *p*-value in the DSC comparison between two-channel and rough mask-only inputs may be due to noise and intensity variations in specific cases, such as Case 19. In this case, variability in the CT image’s pixel histogram likely impacted the performance of the two-channel method, resulting in a decrease in DSC for the single-channel input. This suggests that while the CT image provides valuable context, it can introduce noise in cases with high intensity variation, especially in small datasets. Nonetheless, the two-channel method remains valuable for its comprehensive anatomical insight.

Future work includes: further pre-processing the data to boost model performance, easing our exclusion criteria to train a more general model, externally evaluating the model on other cardiac CT datasets, and replicating our results on other segmentation network architectures (such as the U-Net [6]).

